# Hitchhiking on the Frontier: Accelerating Eusociality and other Improbable Evolutionary Outcomes by Trait Hitchhiking in a Boom-and-Bust Feedback Loop

**DOI:** 10.1101/053819

**Authors:** William H. Calvin

## Abstract

Here I analyze the brush-fire cycle behind the brushy frontier of a grassland, seeking evolutionary feedback loops for large grazing animals and their hominin predators. Several months after a lightning strike, the burn scar grows enough new grass to expand the carrying capacity for grass-specialized herbivores,which evolved from mixed feeders in Africa during the early Pleistocene. The frontier subpopulation of grazers that discovers the auxiliary grassland quickly multiplies,creating a secondary boom for its hominin predators as well. Following this boom, a bust occurs several decades later when the brush returns; it squeezes both prey and predator populations back into the core grassland. This creates a feedback loop that can repeatedly shift the core’s gene frequencies toward those of the frontier subpopulation until fixation occurs. Any brush-relevant allele could benefit from this amplifying feedback loop, so long as its phenotypes concentrate near where fresh resources can suddenly open up, back in the brush. Thus, traits concentrated in the frontier fringe can hitchhike; improved survival is not needed. This is natural selection but utilizing selective reproductive opportunity instead of the usual selective survival. Cooperative nurseries in the brush’s shade should concentrate the alleles favoring eusociality, repeatedly increasing their proportion via trait hitchhiking in the feedback loop.

## Introduction

It is important to analyze evolution’s fast tracks because they can occasionally pre-empt the more familiar slow tracks powered by selective survival. The traditional Darwinian approach looks to some immediate usefulness that allows differential survival to slowly operate on current variations in a trait. Here I am looking instead for an amplifying process in the population ecology, then asking if it could support some form of hitchhiking. A desirable feature of such a process would be amplifying feedback, where some fraction or function of the output feeds back to become part of the input during the next time step, as in the compounding of interest.

The boom-and-bust from cyclical resource fluctuations is a familiar process. On the downside, adaptive evolution can be seen to change gene frequencies via selective survival. A bust also has unusual dynamics in the transition zone between “enough” food and “barely enough,” even if food production is held constant while population grows. Puleston, Tuljapurkar, and Winterhalder (2014) describe what happens when the population size nears the carrying capacity (“The Invisible Cliff”).

Here I address, using a two-compartment model, what can happen on the upside before a boom, how expansion can produce unusually rapid evolution via amplifying feedback during the bust that follows. Not only can allele proportions shift quickly to achieve fixation for a mutation (a “quick fix” indeed) but unrelated traits can be carried along merely by being in the right place at the right time.

### Two rarities seen in human evolution

This analysis of boom-and-bust was prompted by the puzzle presented by two very different evolutionary rarities seen in human evolution.

1. Eusociality, where some individuals compromise their own reproductive success while aiding that of others (as in wet nursing, which reduces fecundity by delaying the return of fertility) is quite rare. Nowak, Tarnita & Wilson (2010) count only 19 instances in all the branches on the animal side of the evolutionary tree where such prosocial behavior has arisen, mostly as sterile castes in insects. The only two mammals known to have eusociality are humans and prairie voles.
2. The three-fold enlargement of our brain over a mere 2.3 million years is also a rarity (Calvin 2017). Its unusual behavioral correlates were singled out early in our understanding of the evolutionary process: that large gap in intellect between great apes and preagricultural humans which so puzzled Alfred Russel Wallace (1869) because he could not imagine a route for selective survival to create this singular enhancement.

Thus two major changes that “make us human” have escaped our Darwinian understanding. A genetic feedback loop provides a new way of approaching each of those rarities. While I have postponed the consideration of brain enlargement per se to Calvin (2017), here I analyze eusociality and the trait hitchhiking common to it and to brain enlargement.

### Non-random gene flow

The fate of a new mutation is known to be affected by a range expansion (Edmonds, Lillie, & Cavalli-Sforza 2004; Excoffier & Ray 2008; Edelaar & Bolnick 2012), achieving fixation more quickly when located just behind an advancing wavefront. Wavefronts have traditionally been analyzed using the diffusion equation or the random walk at an open frontier. Instead, I will be using two thinly connected compartments where the migration time scale is measured in months and there is a feedback loop taking decades to complete.

Edelaar & Bolnick (2012) observe that “Most evolutionary models assume that dispersal is random with respect to genotype.… There is a growing realization that dispersal might not involve the random sample of genotypes as is typically assumed, but instead can be enriched for certain genotypes.” Some hitchhiking is seen at the genetic level, as when genes in the same haplotype move together during crossing over; only one of them is under selection pressure and the others hitchhike.

*Trait hitchhiking,* as I am using the term, instead operates at the level of population ecology where concentrated co-located traits move together during migration. A boom-and-bust feedback loop, as I will show, can pump up the hitchhiking allele’s proportion to fixation in only a few dozen repetitions of the feedback loop. This has implications for the spread of disease as well as the evolutionary aspects of population ecology. I will use two examples to explore the loop’s consequences: the development of antibiotic resistance in bacteria and the evolution of eusociality in humans.

## Conditions for the boom-time

The mile-high savannas of East Africa and South Africa have a high yearly rate of lightning strikes. Many brush fires result and, in the dry season, a large area can burn. Soon, grass sprouts. The grazer population that discovers the fresh grass should quickly double and re-double its population numbers en route to the new carrying capacity, all based on the gene frequencies that characterize the brush fringe with the core grassland. This is an example of non-random gene flow.

In subsequent decades, as returning brush gradually replaces the temporary grass, their offspring are squeezed out of the burn scar (Fig. 3). If they join the parent population, the gene flow changes allele proportions in both the grasslands core and its subsequent frontier subpopulation.

The cycle repeats once another lightning strike sets off the brush-fire feedback loop.Populations of mixed-feeders such as modern elephant and impala need not experience a decades-long change in overall food resources. The leaf-eating browser populations are instead reduced by a brush fire. But for grazers, the auxiliary grassland provides a boom time by extending their range. (Lightning also causes grass fires in the core grassland but grasses recover so quickly that grazing resources are little affected.) Therefore the boom-and-bust feedback loop requires a grazer boom, not a benefit for herbivores in general.

There were surely brush fires older than 2.3 million years ago in the East African Rift Valley but there were no large specialized grazers (Cerling et al 2015), only browsers and mixed feeders that were ineligible for a boom and its feedback loop on bust. Minor climate fluctuations can enhance the amplification process: droughts beforehand or stronger winds make for a larger burn scar, a bigger population boom, more return flow into the core decades later, and thus a larger single-episode boost in the proportion of frontier alleles in the core.

Is this consistent with Darwin’s notion of natural selection?

### The Darwinian process and state-dependent fecundity

For the gradual quality improvement that we associate with natural selection, I earlier identified six essential conditions for a full-fledged Darwinian process (Calvin 1997), which I formulated in more general terms to cover non-genetic examples such as competing cerebral codes during “Get set” movement planning (Calvin 1996). Here I annotate that abstract formulation with examples from gene frequency:

1. There must be a pattern λ involved (such as the ordering of DNAs) that stores information;
2. The pattern λ must be copied somehow (indeed, that which is semi-reliably copied may help to define the pattern).
3. A variant pattern λ′ must arise occasionally (via copying errors, cosmic ray hits).
4. The pattern λ and its variant λ′ must compete with one another for occupation of a limited work space (much as bluegrass and crabgrass compete for space in a back yard).
5. The copying competition between λ and λ′ is biased by a multifaceted environment (for grass: soil moisture, frequency of cropping by herbivores, nutrient availability). This condition is what Darwin meant by natural selection; the preceding four were implied. But these first five are still incomplete; they create a random walk that ignores the whatever current success had been achieved.
6. A variant pattern λ′ is more likely to arise from the more successful of the current patterns λ^1^, λ^2^, λ^3^, simply because the most successful phenotype is more numerous as a target for mutation-making. This last condition is Darwin’s inheritance principle, promoting continuing exploration of the trait’s fit to the phenotype’s environment without losing its history.

Yet there is nothing to prevent a free ride, as when co-located traits “go with the flow” (variously termed transport, bulk flow, and convection, all of the way back to whatever Heraclites called it in ancient Greek 2,500 years ago).

Natural selection operates both by selective survival and by changes in reproductive output (as when food quality improves and double ovulation creates dizygotic sheep twins). While no change in lifetime births per mother (fecundity) is postulated here, the temporary range expansion allows more of her infants to grow up and reproduce themselves^2^. Boom-and-bust feedback loops seem not to require modification of these “six essentials” for present purposes except that Essential 4 now needs to allow the closed work space to occasionally add a closed neighboring compartment (say, the neighbor’s back yard), so that some of the surplus-to-replacement offspring can occasionally survive immaturity.

### Mutations and the gene committee

Here, *new* may be imagined as an allele *λ*′ of a gene that functions within an ensemble of genes *α, β, γ, δ, ε,… λ* controlling the development of some aspect of morphology or behavior.

Functionally, *λ*′ needs to be a small tweak of the usual allele *λ* so that the modified gene can continue to function well enough in the ensemble to be reproduced by a surviving phenotype.

This suggests that the usual mutation of evolutionary significance is a single nucleotide polymorphism (SNP) in the germ line; each human newborn has about 60 new SNPs, ones not seen in either parent (Kong et al 2012), suggesting that each parent accumulated them in the germline prior to their offspring’s conception at the rate of about one per year. Most such newcomers are lost over the generations, though some continue in a balanced polymorphism. However, the feedback loop can allow one acceptable committee newcomer after another to go to fixation. Thus the ensemble *α′, β′, γ′, δ′, ε′,… λ′* may largely consist of “quick fixed” mutations, taken out of competition and no longer able to backslide.

Note that the rate limiter for such evolutionary change could become the mutation rate (as I will argue for the hominin brain enlargement rate; Calvin 2017), rather than the slower speed of the usual selection processes. Mutation rate is not merely the cosmic ray arrival rate; most knockouts are repaired, and so the new SNP rate also reflects the rate of mistakes during gene repair.

### Selective expansion of the frontier allele’s phenotypes

Let *P* represent population numbers, whereas *A* is always a dimensionless proportion for the newest allele λ′ in that population, its “market share.” In the main population *P^core^*, a locus has two alleles, *A^new^* for *λ*′ and *A^old^* for *λ*.

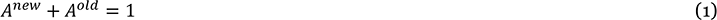

For *A^old^* we can always substitute 1 – *A^new^* or, more simply, use *A* and 1 – *A.*

Near the frontier of the range, allele *λ*′ may be concentrated by the factor *c ≥ 1* because the phenotypes of the original *λ* visit the frontier for briefer periods. By Equation 1, the two frontier proportions are thus:

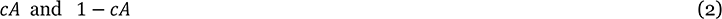

(The actual numbers at the frontier are assumed small and fixed so that the main population 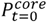 need not be partitioned.)

To count *λ*′ alleles in 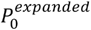, we first define the population’s boom factor *b*

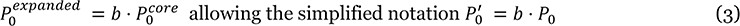

The population in the expansion zone is

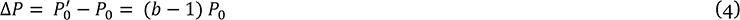

The number of *λ*′ alleles in the original core was *h*⋅*P*_0_*A*_0_; in the surround, there are now 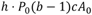 such alleles where the factor 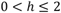 is the number of *A* per individual (for convenience, *h*=1). The number of individuals in the combined population is:

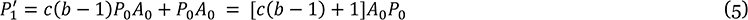

Now we can return to proportions by dividing Equation 5 by the expanded total population 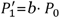 allowing *P*_0_ to cancel out. Thus the allele proportion for *λ*′ at the end of episode 1 is:

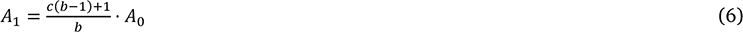

#### The contraction phase

During the bust, one expects the total population 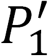 to shrink back to its former value *P*_0_ to be consistent with the core’s prior carrying capacity. For present purposes, I shall assume that the *A*_1_ proportion does not change during downsizing. It is this tweaked proportion *A*_1_ that will be further concentrated by the constant *c* at a future frontier, ready for another boom.

The iterative version of Equation 7 is

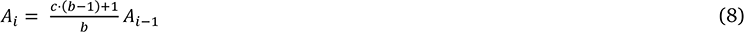

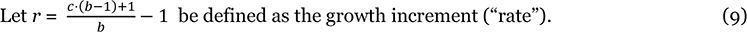

This shows *r* is composed of constants. On the *K^th^* cycle when 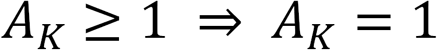 (saturation), the compounded growth of *A* after *K* episodes is:

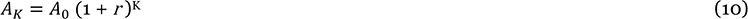

This is the familiar form for the compounding of interest, where the next increment in allele frequency 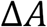 is some fraction *r* of the current base proportion, creating exponential growth in *A* until it fixes at *A_K_* = 1. By the usual evolutionary standards, the growth in *A* toward fixation can be very quick when aided by a feedback loop at the level of population ecology.

The boom-and-bust episodes needed for fixation can be seen in these numerical examples.

##### Example 1.

An initial 20% share and the arbitrary parameters *c* = 1.5 and *b* = 1.3 yield *r* = 0.135 in Equation (9), a 13.5% growth per cycle.

Let *A*_0_ = 0.2 before the first feedback episode.

*A*_1_ = 0.2 (1.135)^1^ = 0.227

*A*_2_ = 0.2 (1.135)^2^ = 0.258

… *A*_13_ ≥ 1 (*k* = 10 is where slowing starts because *c* drops to 1.)

Note that saturation will first occur in the frontier zone at loop *k*, after which the main’s amplification slows somewhat as the cycle increment thereafter remains constant until cycle *K* when the core goes to fixation as well.

##### Example 2.

In Equation (9), if *c* = 1.20 and *b* = 1.3, yielding *r* = 0.046, it results in a more modest 4.6% growth rate,

*A*_1_ = 0.2(1.04615)^1^ = 0.2092

*A*_2_ = 0.2(1.04615)^2^ = 0.2189

… *A*_36_ ≥ 1
and thus *new* has become the only allele at this locus by the 36th cycle, doubly present in all individuals (“fixed”) and so taken out of competition until another SNP succeeds in becoming a functionally^3^ different allele.

### Leak limitations in a center-surround geometry

One expects the borders of the species range to concentrate some traits such as robustness, if only because those lacking the trait promptly circulate back into the central population rather than lingering. After slow expansions in range and many generations, the gene frequencies of the central population should become more like those of the frontier’s gene pool. Leaks, however, reduce the loop’s effectiveness.

- When resources expand quickly, as does grass after a brush fire, some of the core population can leak past the frontier’s concentration of *A* en route to the new resources. This is a diminution of the step up in *A* for the expansion. But there is also a continuing problem.
- The center-surround boom-and-bust lacks any multiple-generation isolation of the surround zone’s subpopulation to enforce inbreeding. The leaks continue but the concentrating does not, and so the inbreeding that helped to maintain *cA* in the frontier zone is no longer as effective during the decades until the bust-phase contraction enriches *A* in the core.

For rapid expansions, a two-compartment model with a narrow passage yields better insights. It also better fits the geometry for the boom-and-bust seen in large herbivores specialized for grass (in the East African Rift Valley, the present-day grazers are wildebeest and zebra). The two-compartment model’s gateway corridor (seen in Fig. 2) better limits the continuing dilution before the bust enriches *A* in the core.

**Fig. 1.**
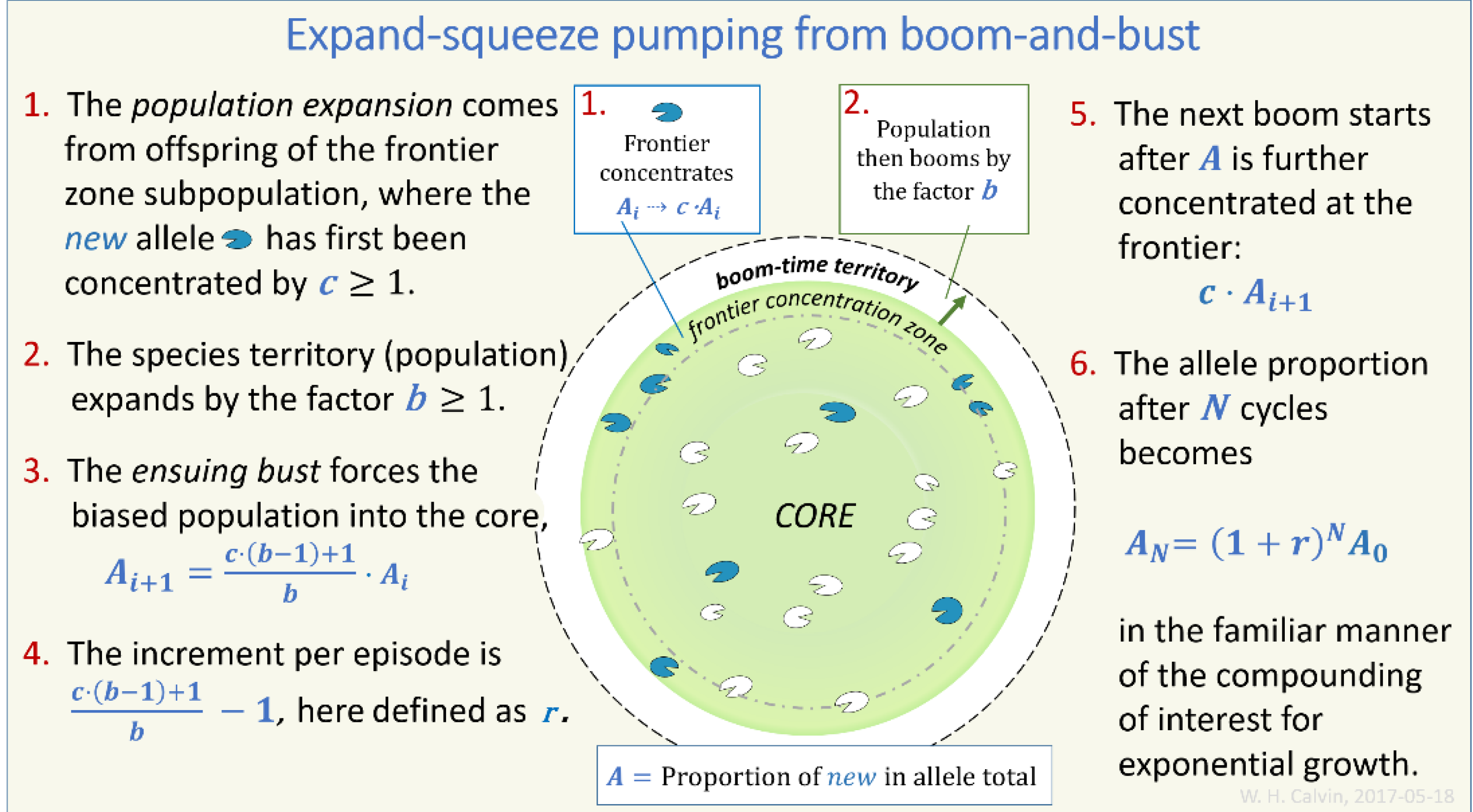
Expand-squeeze pumping. The center-surround geometry and the compounding of allele proportions.

**Fig. 2.**
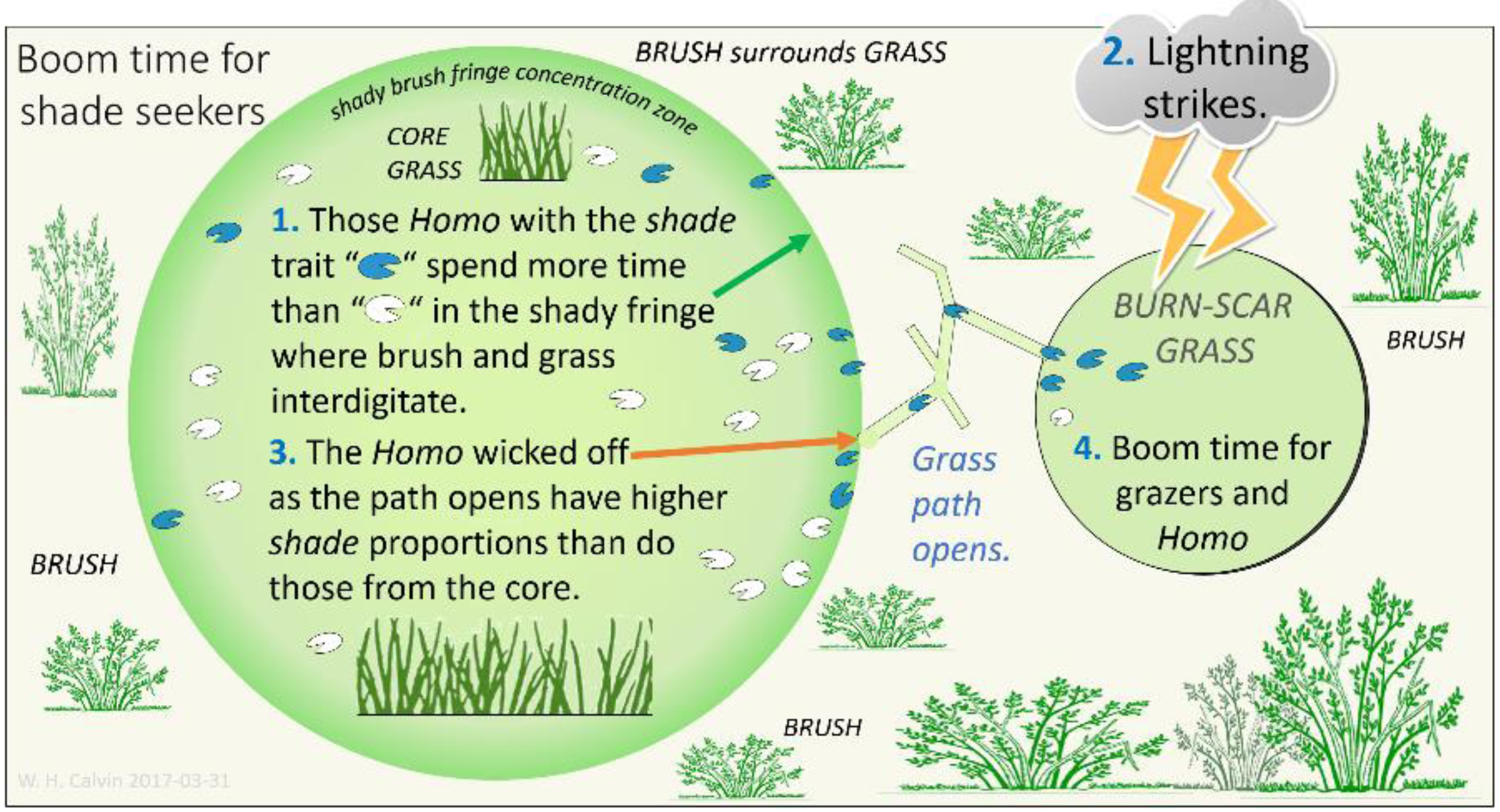
Boom time for a concentrated allele. In ordinary times, those with the *shade* trait are likely to spend more time in the brush fringe before leaving. This enrichment of *shade* in the fringe means that when a path opens to the burn-scar’s temporary grassland, those wicked off into it are not those with the usual allele proportions of the core.

## Why brush matters for trait hitchhiking

In Fig. 2, the boundary of the grass resource in both compartments is brush, not the absence of grass as in Fig. 1. There is a frontier fringe where brush and grass interdigitate (Calvin 2017), forming dead-end paths back into the brush. Lightning strikes farther back in the brush and, in the dry season, a large burn scar will develop within a week. The brush-fire scar turns into a temporary grassland within a month or two.

The more cautious herbivores often avoid brush paths as predators can hide there. They pack together in grassy patches within the brush, the reason why bush pilots must first scare the herbivores off the landing strip before making a second pass to touch down. Yet the dry season may send even the cautious grazers deeply into brush’s byways.

The burn-scar grass may be discovered via the usual process of trampling new pathways through the brush fringe during the dry season. Aided by the excrement left behind, encroaching brush may close the path the next year. However, some paths turn out not to be “dead end” after all. This range expansion into the auxiliary grassland via a narrow corridor (Fig. 2) now supports a total population *P*′ but the second compartment has the concentrated allele proportions of the old frontier fringe, not those for the core grassland population.

Equation 6 still applies, as the constants *b* and *c* are not dependent on the center-surround geometry of Fig. 1 Indeed, because of the unstated dilution problem in center-surround pumping, the pathway to the auxiliary grassland provides a far better illustration of Equation 6; the pathway may even close a year after opening because the fertilization of leafy plants creaes growth that obscures it.

As before, brush eventually returns to the burn scar (its groundwater supply is not reduced by the burn, so leafy “volunteers” amid grass now survive the dry season). Fig. 3 illustrates the squeeze.

**Fig. 3.**
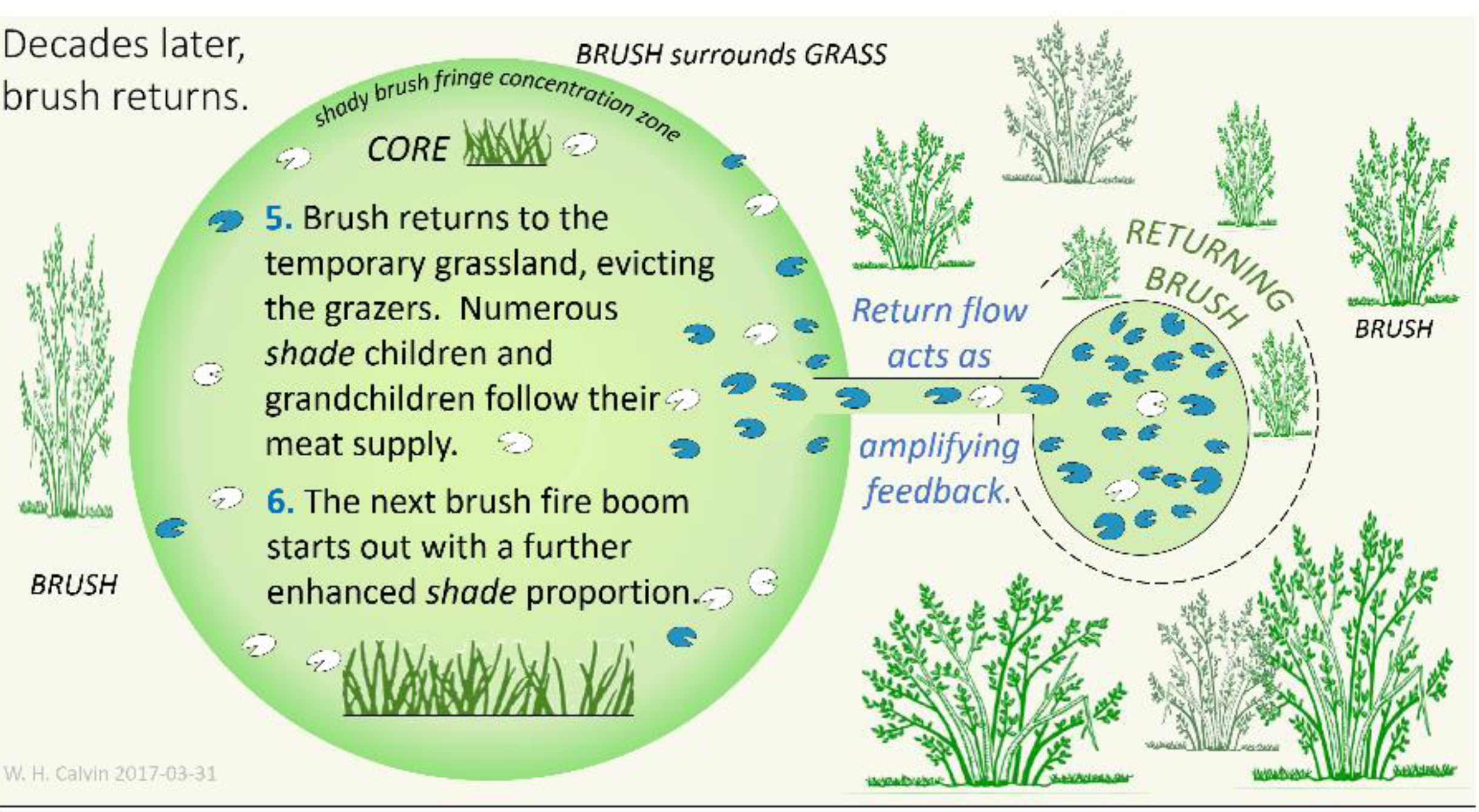
The bust and the feedback it creates. Decades later, those leafy volunteers in the temporary grassland are finally taking over, suppressing grass growth nearby. As grazers and their predators are pushed out, the core population’s allele proportions are enriched by the arriving shade phenotypes.

### Shade-seeking traits can hitchhike in the feedback loop

By hanging out in frontier zone that benefits from the occasional boom in the meat supply, hitchhiking is enabled for traits among the meat-eaters that tend to concentrate in the frontier fringe, often for reasons other than acquiring meat. Again, this is habitat preference via dwell time: some bearing the allele *λ*′ simply stay longer in the fringe before circulating back out into open country that lacks shade.

Hitchhiking traits prosper not by their own usefulness but because their phenotype got a free ride to a population boom by hanging out in the frontier fringe, which gives them sudden access to a burn scar upon occasion. Traits such as food preference and shade-seeking habitat preference are obvious candidates for promoting prolonged dwell time and, decades later, core enrichment for their alleles.

I will use *shade* in place of *new* or *λ*′ because the hitchhiking traits I will discuss all involve shade-seeking habitat preference. In *Homo*, shade-seeking in brush could coevolve with such traits as toolmaking, food preparation, fire-starting, and enhancing the communal rearing aspect of eusociality. The boom and its feedback loop would then promote genes for those hitchhiking traits, once the bust occurs and amplification feedback is created by spreading them into the core. Around the loop, there need not be a filter via selective survival, which is merely one way that relevant alleles can be concentrated in the frontier zone.

Most obviously, the brush fire loop should amplify habitat preference alleles for the frontier, even in the central core population that rarely visits the frontier. But it also amplifies any trait that co-locates in the brush, provided it has a concentration gradient between the core and the periphery; their phenotypes also get the benefit of having more surplus-to-replacement offspring surviving to reproductive age.

## An abscess’s autocatalytic alternative

The basic components of the allele-promoting feedback loop, stated more abstractly, would seem to be

1. an allele concentration mechanism in an opportunity zone,
2. a population that can boom out of the opportunity zone into unused resources, and
3. a migration path from the boom territory back into the core population during the bust phase, needed to create the feedback loop.

One can use these three features as a search image in prospecting for other evolutionary feedback loops in nature. But a further simplification can be seen in briefly considering the following situation from medicine, which initially looks nothing like the brush-fire geometry.

An abscess in weight-bearing skin (“bed sore”) is thought to arise from impaired circulation. The smaller arterioles and capillaries serving the abscess can be collapsed by the body’s weight, backing up blood in proximal arteries until the weight is relieved once an hour by rolling over the patient. This leads to a long residence time for the trapped blood in vessels that are narrow enough to distort the shape of the red blood cells. This tight fit constitutes a concentration zone: antibodies and antibiotics have a long time in which to act on the vessel’s content, whose cells cannot just bounce away as they do in the general circulation. Those bacteria with an allele that confers antibiotic resistance would gain “market share” as the vulnerable alternatives swell and burst as antibiotics cause the bacterial cell to retain water.

Let us suppose that freely-circulating blood has bacteria, of which 1% are antibiotic resistant. In the weight-bearing skin where blood in narrow vessels is stopped, let us suppose that antibiotics kill half of the ordinary bacteria. Their removal serves to increase the resistant bacteria’s proportion to 50%. This is the blood that, once passed through the abscess, is returned to the central venous circulation to create amplifying feedback; there is of course an enormous dilution by blood returning from tissues without a circulation slowdown, so the original 1% of resistant organisms may merely be increased to ≈ 1.01%. But this feedback loop continues to boost the percentage with each flushing of the line, 24 times a day.

In this simplified analysis, nothing depends on what the abscess itself contributes. Indeed, we can do without an abscess at all and the periodically backed-up blood vessels should still promote antibiotic resistance. Perhaps an abscess forms to sequester the decomposing bacteria arriving from the arterial side so that they do not clog up the venous passage and cause tissue necrosis.

This account has variants on all three features suggested for the search image:

a. an allele concentration mechanism (selective survival in backed-up small blood vessels),
b. pulse to flush the line (those hourly rollovers function much like the bust), and
c. migration path back into the core population (the general circulation), creating the feedback loop (making antibiotic resistance more and more common in the general circulation).

Thus this antibiotic resistance allele pump does not require a boom per se, though boom-and-bust can provide larger changes per episode.

This invites an analogy to the autocatalytic processes in chemistry. An ordinary catalyst features an enclave where entering molecules may rattle around inside, losing their momentum until it is little different from that of other trapped molecules. Collisions may then cause some molecules to stick, rather than ricochet, forming chemical bonds at a much faster rate than matched-momentum collisions in the open.

But an autocatalytic process is defined for a chain of chemical reactions that, once started, maintains itself at some rate, usually because the last of the chain produces a byproduct that the initial reaction can use as fuel. It does not require a continuing train of triggers such as booms and rollovers. Yet concentrating an allele in the loops’ opportunity zones itself can be considered autocatalytic in the sense that more *shade* individuals from the core will now feel an urge to seek out shade, increasing the number of visits to the brush fringe frontier beyond what it is from random walks alone, a “pre-treatment” to the further concentration of the *shade* allele in the frontier zone. Now the *shade* allele not only stays for longer visits but they increasingly make more visits.

## Eusociality is promoted by a frontier’s shade

My second example of an allele pump involves trait hitchhiking for prosocial behaviors in the context of the basic brush-fire feedback loop for grazers and their followers. Eusociality is rarely seen in evolutionary lineages (Nowak, Tarnita & Wilson 2010); there are only two examples among mammals and one is in the *Homo* lineage.

Eusociality is where some individuals reduce some of their own lifetime reproductive potential to raise the offspring of others, underlies the most elaborate forms of social organization seen in evolution. As noted, breast feeding someone else’s infant serves to suppress ovulation in the nurse, prolonging lactational amenorrhea and reducing her fecundity. Might eusocial variants concentrate in the frontier fringe, their allele later amplified in the core by the same biased-boom-and-return loop that affects the grazing animals and feeds their predators?

In addition to the tendency of many animals to stay out of the midday sun, human infants may need sun protection all day because their small bodies have a lot of bare surface area for their volume, which causes them to gain heat more quickly than their parents. Infants can suffer heat stroke on hot days when they are not being held against a large heat sink supplied by blood that has been evaporatively cooled elsewhere. While cuddling an infant serves to warm it when it is chilled, source and sink can be reversed on hot days.

Before the sling was invented to hold an infant beneath a breast, a mother needing to work with both hands required another large-enough individual to cuddle the infant. Alternatively, this working mother could have parked the infant in the shade where someone could monitor it, perhaps even nursing it. Shade thus serves to concentrate community nursery traits in the frontier fringe, where they occasionally benefit from the enhanced grand fecundity afforded by a burn scar’s population boom.

The repeated booms over many generations could keep shifting the overall population toward eusociality, even if there is no selective survival judging its usefulness, simply because co-located traits can boom together during the resource expansion. In contrast, the frontier’s selective survival for robustness is slow to alter the genetic makeup of the core by continuous diffusion because of the large numerical disproportion between the frontier rim and the core population.

A feedback loop, however, squeezes numerous brush frontier genes for eusociality into the core, based merely on who was in the right place (the frontier fringe) at the right time (when the temporary grassland opened up). Many other frontier-concentrated alleles (for such traits as toolmaking, food preparation, and fire-starting) could be amplified if they were to maintain an allele concentration gradient between the core and the frontier (Calvin 2017).

## Discussion

The feedback loop provides more than the evolutionary overdrive that one might expect from an analogy to catalysts. Trait hitchhiking better resembles an individual’s free ride up an escalator toward increased offspring survival, where a habitat preference for the escalator’s entry location enables this exception to the familiar process of changing morphology and behavior via selective survival of variations. Because a variant may be quickly driven to fixation, the feedback loop boosts the speed of evolution.

The other boost is for the evolutionary creation of a novel solution. Trait hitchhiking joins succession of function (Dohrn 1875, Caianiello 2015) as an example of how a novel function can be promoted. Darwin’s example (1859) of conversion of function was the fish’s swim bladder, whose gas exchange with blood is used for regulating buoyancy via an expanding air sack. This bladder was converted into an organ specializing in gas exchange between blood and inhaled air.

The head start provided by an existing adaptation sometimes promotes an elaborate secondary use, one that itself has no history of demonstrated usefulness (Calvin 2004). The feedback loops examined here suggest that seeming discontinuities in evolution may have a long “pre-adaptation” phase at little cost before the evolutionary payoff.

> I thank my University of Washington colleagues James J. Anderson, Katherine Graubard, and Charles D. Laird for discussions on the manuscript and numerous CARTA colleagues for the genetics education.

## Footnotes

1

2 Boom-time amplification is consistent with a generation-skipping definition we might call “grand-fecundity.” Counting *grandchildren* per *grandmother* rather than lifetime *children* per mother can allow for environmental influences that are state-dependent, such as a boom time that temporarily reduces the usual immature mortality. Fecundity is already based on surviving the first nine months and it is now known there are state-dependent environmental influences on the survival of human embryos. Some drinking-water sources promote a high rate of spontaneous abortion: sometimes more than half drop out (Swan et al 1998) before the heartbeat begins six weeks after conception in humans; there is a 10-15% “miscarriage” rate thereafter. Assuming that the lowest dropout rate seen, 27%, represents the baseline from zygote developmental misfits, one must ask if the more elevated dropout rates such as 70% involve the selective elimination of some genotypes to produce a hidden environmental bias of the characteristics of the newborn population.

3 Some SNP mutations merely code for one of the other nucleotide triplets specifying the same amino acid. For example, if you start with the usual codon *GCU*, you get the amino acid Alanine tacked on the lengthening string of amino acids which will fold up to become a protein influencing some function. If the SNP of *GCU* produces *GCC*, you still get Alanine. The same for *GCA* and *GCG*. Function does not change, at least in the present generation. But if the following generation started with *GCC* in the germ line instead, a SNP of it might be the *ACC* codon, and so a different amino acid, Threonine, is added to the chain instead of Alanine. This may change the way in which the nascent protein can bend and fold. In this example, only *ACC* qualifies as a new allele: it alone changes the protein’s function (facilitating prostate cancer in this case).

## References

Caianiello S (2015) Succession of function, from Darwin to Dohrn. Hist Philos Life Sci. 36: 335–345.10.1007/s40656-014-0041-y

Calvin WH (1996) The Cerebral Code. Cambridge MA: MIT Press.

Calvin WH (1997) The six essentials? Minimal requirements for the Darwinian bootstrapping of quality. Journal of Memetics 1:1, at http://cfpm.org/jom-emit/1997/vol1/calvin_wh.html

Calvin WH (2004) A brief history of the mind. New York: Oxford University Press

Calvin WH (2017) In Rift Valley settings with a feedback loop, assortative mating for versatility predicts hominin brain enlargement in some detail. Preprint at bioarxiv, doi: 10.1101/053827.

Cerling TE, Andanje SA, Blumenthal SA, Brown FH, Chritz KL, Harris JM, et al (2015) Dietary changes of large herbivores in the Turkana Basin, Kenya from 4 to 1 Ma. Proc Natl Acad Sci USA. 112: 11467–11472.doi:10.1073/pnas.1513075112

Darwin C (1859) On the origin of species. 3^rd^ edition. London: Murray

Dohrn A (1875) Der Ursprung der Wirbelthiere und das Princip des Functionswechsels: genealogische Skizzen. Leipzig: Engelmann

Edelaar P, Bolnick DI (2012) Non-random gene flow: an underappreciated force in evolution and ecology. Trends Ecol Evol 27: 659–665.10.1016/j.tree.2012.07.009

Edmonds CA, Lillie A, Cavalli-Sforza L (2004) Mutations arising in the wave front of an expanding population. Proc Natl Acad Sci USA, 101(4): 975–979, 10.1073/pnas.0308064100

Excoffier L, Ray N (2008) Surfing during population expansions promotes genetic revolutions and structuration. Trends Ecol Evol 23:34–351.

Kong A, Frigge ML, Masson G, et al (2012) Rate of de novo mutations and the importance of father’s age to disease risk. Nature 488: 471–475 (2012), 10.1038/nature11396

Nowak MA, Tarnita CE, Wilson EO (2010) The evolution of eusociality. Nature 466:1057–1062. 10.1038/nature11396

Puleston C, Tuljapurkar S, Winterhalder B (2014) The invisible cliff: abrupt imposition of Malthusian equilibrium in a natural-fertility, agrarian society. PLoS ONE 10.1371/journal.pone.0087541

Swan SH, Waller K, Hopkins B, Windham G, Fenster L, Schaefer C, Neutra RR (1998) A prospective study of spontaneous abortion: relation to amount and source of drinking water consumed in early pregnancy. Epidemiology 9: 126–133.

Wallace AR (1869) The Malay Archipelago: the land of the orang-utan and the bird of paradise; a narrative of travel, with studies of man and nature. London: Macmillan.

